# Extracting neural signals from semi-immobilized animals with deformable non-negative matrix factorization

**DOI:** 10.1101/2020.07.07.192120

**Authors:** Amin Nejatbakhsh, Erdem Varol, Eviatar Yemini, Vivek Venkatachalam, Albert Lin, Aravinthan D.T. Samuel, Liam Paninski

## Abstract

Extracting calcium traces from populations of neurons is a critical step in the study of the large-scale neural dynamics that govern behavior. Accurate activity extraction requires the correction of motion and movement-induced deformations as well as demixing of signals that may overlap spatially due to limitations in optical resolution. Traditionally, non-negative matrix factorization (NMF) methods have been successful in demixing and denoising cellular calcium activity in relatively motionless or pre-registered videos. However, standard NMF methods fail in animals undergoing significant non-rigid motion; similarly, standard image registration methods based on template matching can fail when large changes in activity lead to mismatches with the image template. To address these issues simultaneously, we introduce a deformable non-negative matrix factorization (dNMF) framework that jointly optimizes registration with signal demixing. On simulated data and real semi-immobilized *C. elegans* microscopy videos, dNMF outperforms traditional demixing methods that account for motion and demixing separately. Finally, following the extraction of neural traces from multiple imaging experiments, we develop a quantile regression time-series normalization technique to account for varying neural signal intensity baselines across different animals or different imaging setups. Open source code implementing this pipeline is available at https://github.com/amin-nejat/dNMF.

## 1 Introduction

Recent advances in imaging techniques have enabled the capture of functional neural ensembles *in vivo* within a wide variety of animal models [Flusberg et al., 2008, Ahrens et al., 2013, Prevedel et al., 2014, Mann et al., 2017]. Demixing the recorded video signals into estimates of individual neural activity remains a critical bottleneck in the analysis of these large and complex datasets. Previous approaches for extracting individual neural activity traces have involved either region of interest (ROI) methods [Kerr et al., 2005, Niell and Smith, 2005, Dombeck et al., 2007, Göbel et al., 2007, Tian et al., 2009, Kerlin et al., 2010, Hofer et al., 2011, Venkatachalam et al., 2016b, Nguyen et al., 2016, Barbera et al., 2016] or matrix factorization methods based on principal components analysis (PCA) or independent components analysis (ICA) [Stetter et al., 2001, Siegel et al., 2007, Reidl et al., 2007, Mukamel et al., 2009] or sparse coding [Pachitariu et al., 2013, Pachitariu et al., 2017].

Non-negative matrix factorization (NMF) [Paatero and Tapper, 1994, Lee and Seung, 1999, Lee and Seung, 2001] based models have been introduced to demix signals from recordings of calcium activity [Andilla and Hamprecht, 2013, Haeffele et al., 2014, Andilla and Hamprecht, 2014, Maruyama et al., 2014, Pnevmatikakis et al., 2016, Pachitariu et al., 2017, Zhou et al., 2018]. A prerequisite for the success of these methods, to permit blind-source separation, is that the imaged ROI remains motionless even when the animal is awake, satisfying the assumption that the spatial footprints of signal sources remain stationary. To facilitate NMF assumptions and remove excess motion variability, a common pre-processing step before NMF is the registration of the imaging volumes to a common template space.

There is a wealth of literature in the medical imaging community regarding the registration of volumetric images to template volumes to account for morphological variability [Klein et al., 2009]. These methods have proven to be very effective in registering images that have similar intensity profiles but they tend to introduce artifacts when the template image and the moving image have different appearances, low signal to noise ratio, or abnormalities [Zeng et al., 2016]. Furthermore, the computational complexity of these methods is a bottleneck since there are potentially tens of thousands of frames in volumetric calcium videos that need to be registered. A number of pipelines [Dubbs et al., 2016, Pnevmatikakis and Giovannucci, 2017, Pachitariu et al., 2017] implement existing sub-pixel registration techniques [Guizar-Sicairos et al., 2008] to enable the rigid and non-rigid registration of calcium videos in a computationally efficient manner. Assuming that the motion does not involve large shifts in the field of view (FOV), these techniques aim to register individual video frames to a template frame through fast patchwise rigid transformations. However, they too are not built to handle severe deformations and large intensity variations.

Recent whole-brain imaging techniques of the model organism *C. elegans* [Schrödel et al., 2013, Prevedel et al., 2014, Kato et al., 2015, Nguyen et al., 2016, Venkatachalam et al., 2016b] have opened up an exciting new avenue of research, enabling simultaneous recording of neural dynamics and freely-moving behaviors in the same animal. Even during restrained imaging, worms can exhibit highly-nonlinear motion [Girard et al., 2007, Larsch et al., 2013, Voleti et al., 2019], violating the assumptions that enable NMF-based signal separation and overstretching the capabilities of fast piecewise rigid registration techniques. Therefore, common approaches have been to apply motion tracking and simple pixel-averaging around cellular tracking ROIs in two discrete steps, often followed by time-consuming supervision and manual correction of the results [Kato et al., 2015, Venkatachalam et al., 2016a, Nguyen et al., 2016]. One way to perform motion tracking is to use a second imaging channel to record a temporally-invariant fluorescent marker (such as RFP) which is insensitive to calcium activity. By using such cellular motion tracking markers, calcium activity can then be extracted by averaging the pixel values in the ROI that overlap with the marker. However, this approach is flawed for at least two reasons: 1) ROI averaging in densely-packed cell regions is prone to mixing signal between different neurons, due to limitations in optical resolution, and 2) introducing a second imaging channel effectively requires experimenters to reduce the frame rate and/or spatial resolution by at least half in order to acquire this channel or add an additional optical path and camera. On the other hand, if tracking is performed only on the calcium imaging channel, due to the low signal-to-noise regime and calcium signal fluctuations, tracking approaches may miss cellular markers at time points when the cells become dim, creating downstream errors in tracking and demixing.

In general, tracking cells in moving animals (and even restrained animals with restricted mobility), has proven to be a challenging machine vision problem [Hirose et al., 2017]. Cell nuclei have similar shapes, thus providing only a limited set of unique features to facilitate their tracking. Spatial noise represents a further, inherent limitation, due to the microscopic size of the objects under investigation. Most available microscopy approaches scan the animal in both space and time to achieve volumetric video recordings. Therefore, there are fundamental limits in reaching the high spatiotemporal resolution necessary to resolve unique cell identities and extract their calcium signals through tracking techniques.

Even if high accuracy cell tracking can be achieved, another issue with extracting calcium signal around tracked ROIs is that many existing volumetric optical imaging setups have a relatively poor resolution in the depth axis, characterized by an elongated point-spread-function [Yang and Yuste, 2017]. This phenomenon causes the calcium signals of nearby cells to be mixed, which in turn causes the pixel-wise signal read-out to be an inaccurate portrayal of actual neural activity.

Orthogonally, there have been NMF techniques that are invariant to signal shift, such as convolutive NMF [O’grady and Pearlmutter, 2006, Smaragdis, 2006]. However, these techniques model discrete translation based shifts and are not suitable for modeling the complex deformable motion exhibited across biological volumetric recordings.

In the case of *C. elegans* imaging, worms can exhibit nonlinear motion (even when immobilized using popular paralytics [Larsch et al., 2013, Venkatachalam et al., 2016a, Voleti et al., 2019]) and variability in their neural firing patterns over time, making the application of previous techniques such as Normcorre [Pnevmatikakis and Giovannucci, 2017] or convolutive NMF ineffective. To surmount these issues, we introduce deformable non-negative matrix factorization (dNMF) to jointly model the motion, spatial shapes, and temporal traces of the observed neurons in a tri-factorization framework. Instead of the two-step approach of sequentially tracking then demixing calcium signals, we update motion parameters together with updates in the spatial and temporal matrices. To ensure that our model is not overfitting and picking up spurious motion and signal, we use regularized models for cell shapes, temporal fluctuations, and deformations. The model parameters capture the worm’s motion corresponding to a fixed, spatial representation of the video, enabling the deformation terms to match the worm’s posture at each time frame. Our framework is general and is suitable for decomposing videos into a set of motion parameters, fixed spatial representations for image components, and temporally varying signals with underlying linear and/or nonlinear motion. This approach can be considered a generalization of the model developed in [Peng et al., 2012] (applied to calcium imaging data by [Poole et al., 2015]), which restricts attention to affine transformations.

We validate our method on an intensity-varying particle-tracking simulation and compare it to state-of-the-art calcium-imaging motion-correction techniques [Pnevmatikakis and Giovannucci, 2017] followed by NMF [Pnevmatikakis et al., 2016]. We then demonstrate the ability of our framework to extract calcium traces from all neurons in the head and tail of semi-immobilized *C. elegans* exhibiting nonlinear motion. We use a dataset of 42 animals, 21 worm heads and 21 worm tails, recorded for 4 minutes each while presenting three stimuli, a repulsive concentration of salt and two attractive odors. We find that the proposed approach outperforms both ROI averaging and standard NMF, delivering more accurate tracking and demixing than either of these methods in this dataset.

Finally, after accurate extraction of neural activity signals from each animal, a post-processing normalization step is still required in order to compare neurons of the same type, across a population of animals. This is because factors such as variable illumination, anisotropy associated with animal orientation, and a lack of stereotypy in fluorescence expression across animals introduce substantial variability into the baseline and amplitude of the extracted neural activity signals. Standard post-processing approaches based on estimating Δ*F*/*F*_0_, do not resolve this variability, which if uncorrected will confound any group-type neural comparisons across animals. Even worse, outlier signals that arise due to mistracking and demixing can considerably warp the mean signal measured across a population of animals, especially when the neuron type of interest is dim and is present next to neuron types with brighter signal.

To reduce this excess variability across animals, we introduce a time-series normalization approach, termed quantile regression. This approach optimizes for a linear transformation of time-series intensities in a group of samples (e.g., all traces extracted from a given cell type over all animals), transforming the time-series samples to have matched histograms. We compare this approach with z-scoring and advocate its adoption for population-based time-series analysis due to several desirable properties. In particular, our approach retains the approximate baseline and magnitude across a population of neurons of the same type, while maintaining robustness against outlier signals. Lastly, we introduce an option for ensuring the non-negativity of the normalized signals, when appropriate for the biological measurements being performed.

## 2 Methods

The joint motion correction and signal extraction framework proposed here involves several steps illustrated in Figure 1. First, the volumes undergo several pre-processing steps that involve coarse tracking, background subtraction and smoothing, details of which are discussed in the "Pre-processing steps" subsection below. The pre-processed volumes are then subjected to simultaneous deformation compensation and signal demixing using a matrix tri-factorization model.

**Fig. 1.**
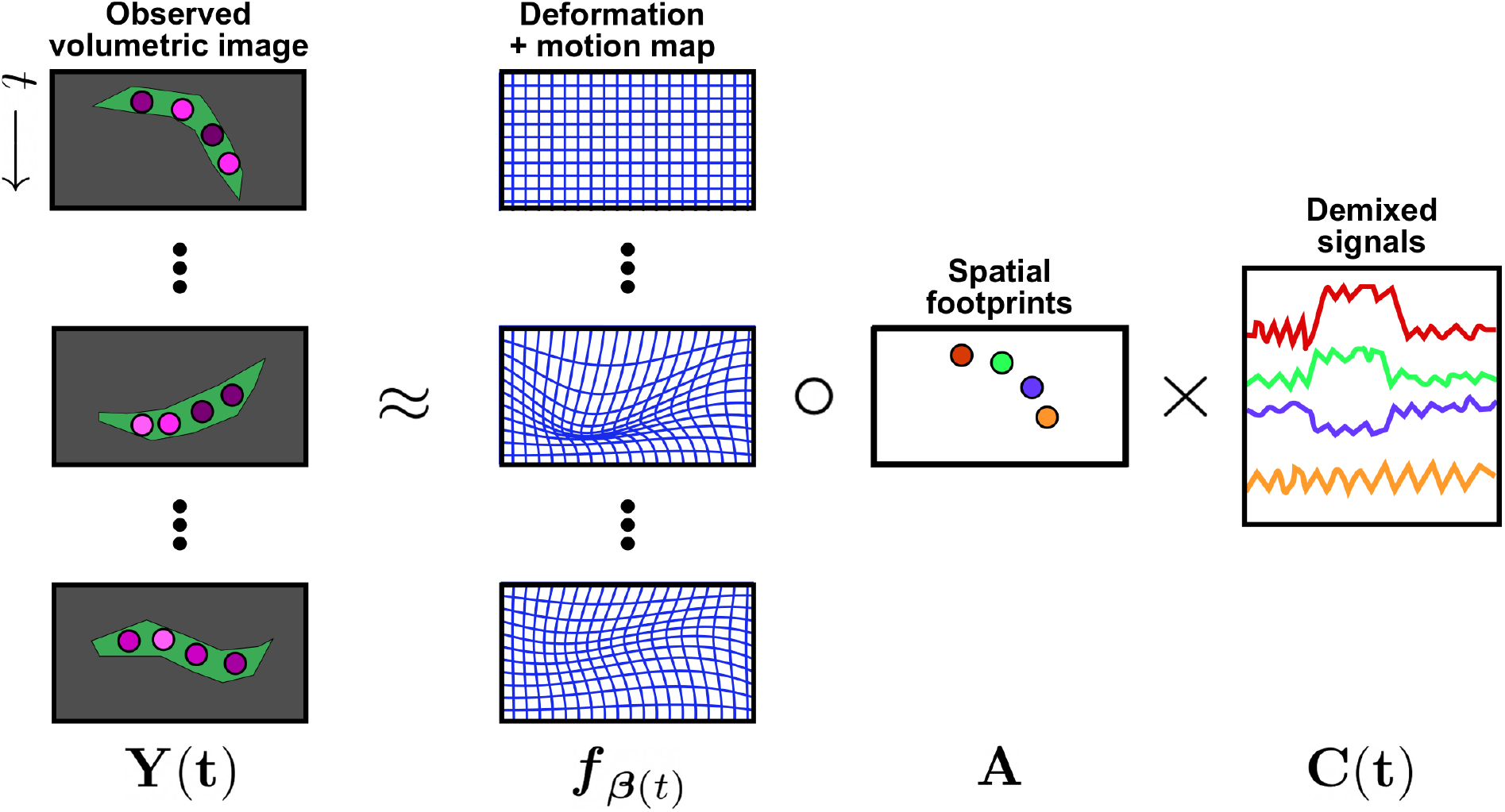
Schematic of the deformable non-negative matrix factorization model. The volumetric time series data ***Y*** (*t*) is factorized into time-varying deformation + motion maps ***f***_β(*t*)_ which transform the factorized signal (with spatial footprints ***A*** multiplied by time-varying intensity coefficients, ***C***(*t*)) onto the observed data volumes.

First, we introduce notation. Let ***Y***_*t*_ ∈ ℝ^*d*^ denote the *d*-pixel vectorized volumetric image at time *t* = 1,…,*T*. We seek to decompose the observations, ***Y***_*t*_, into a factorization involving a time-varying deformation term, 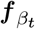 acts on a time-invariant canonical representation of *k* shapes encoded by ***A***. The time-varying spatial signatures, 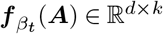 are then multiplied by signal carrying coefficients ***C**_t_* ∈ ℝ^*k*^. We also encourage model parameters to be “well-behaved” using regularization functions, 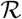 (details of which will be outlined later). The resulting objective function is:

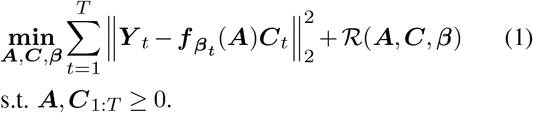

This formulation differs from standard NMF techniques [Lee and Seung, 2001] in that the spatial footprint term consists of a time invariant term, ***A*** and a time varying term, 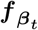, which is a differentiable transformation parametrized by β_*t*_, that deforms the canonical representation into the *t*-th time frame. β_*t*_ encapsulates the motion parameters and is usually low dimensional to avoid over-parameterization and overfitting. The regularization 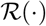 further constrains the possible choice of spatial footprints, signal coefficients, and spatial deformations. Figure 1 illustrates the model. Next, we detail two possible parameterizations of the spatial terms, ***A*** and ***f***.

### 2.1 Spatial component: non-parametric model

Similar to the standard NMF models, we can parameterize ***A*** using a *d*-by-*k* matrix, where *d* is the number of pixels of one time frame of the video and *k* is the number of objects that are present. We use a Gaussian interpolant, ***T**_t_*, to transform these spatial footprints to arbitrary locations such that 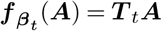, where ***T**_t_*: ℝ^*d*×*d*^ and

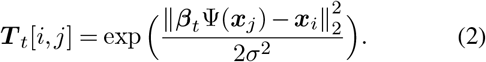

Here, ***x_i_***, ***x_j_*** ∈ ℝ^3^ denote the coordinates of two arbitrary pixels in the volume. Ψ: ℝ^3^ → ℝ^*p*^ denotes a basis mapping of coordinates to enable non-linear deformations and ***β**_t_* is a 3-by-*p* matrix that parametrize the deformations. For example, in the case of a quadratic polynomial basis, ***β**_t_* would be a 3-by-10 matrix, and Ψ: ℝ^3^ → ℝ^10^ would be the quadratic basis function Ψ([*x, y, z*]^*T*^) = [1*, x, y, z, x*^2^, *y*^2^, *z*^2^, *xy, yz, xz*]^*T*^. The choice of *σ* controls the amount of the spread of the mass of a pixel into nearby pixels. Further details on optimization in this model can be found in the appendix 1.

### 2.2 Spatial component parametrization: Gaussian functions

When we have strong prior information about the component shapes we can incorporate that into the model using an appropriate parameterization for the spatial footprints. Neural activity is most commonly imaged using cytosolic or nuclear-localized calcium indicators; nuclear-localized indicators can be reasonably modelled using ellipsoidally-symmetric shape models. Specifically, we observed that the spatial component of the neurons in the videos analyzed here, of *C. elegans* imaged using nuclear-localized calcium indicators, can be well approximated using three-dimensional Gaussian functions. By taking advantage of this observation we can reduce the number of parameters in ***A*** from one parameter per pixel per component, to *k* 3D centers (3 parameters per each neuron) and *k* covariance matrices (6 parameters per each neuron using the Cholesky parameterization). Formally, we model the footprint of component *k* using a 3-dimensional Gaussian function with location parameters ***μ***_*k*_ ∈ ℝ^3^ and shape parameters **Σ**_*k*_ ∈ ℝ^3 × 3^. Under this new spatial model for ***A*** = {***μ***_1:*K*_, **Σ**_1:*K*_}, we modify the 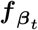 function to match this parameterization to have 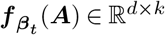:

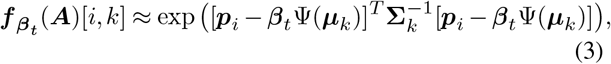

where ***p**_i_* is the 3D coordinate of the *i*-th pixel in the image. (Note that non-negativity of the spatial components is enforced automatically here.) Due to the differentiability of 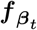, it is straightforward to compute gradients with respect to ***β**_t_* and **Σ**_*k*_.

### 2.3 Regularization: temporal continuity

To enforce smoothness of the temporal traces and motion trajectories in time we add a regularizer that penalizes discontinuities in the neural trajectories and signal coefficients. Specifically, we encourage the neural centers and signal coefficients at neighboring time points to be close. The regularizer for this purpose is:

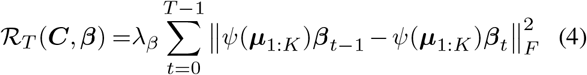

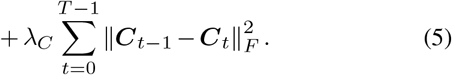

In this formulation *ψ*(***μ***_1:*K*_) is the quadratic transformation of the canonical neural centers. When multiplied by ***β***_*t* - 1_ and ***β***_*t*_ the result will be the neural centers at time *t* - 1 and *t* respectively.

### 2.4 Regularization: Jacobian constraints for plausible deformations

The term 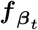 induces a deformable transformation of the pixel correspondences between time *t* and the canonical representation ***A***. In order to constrain this transformation to yield physically realistic deformations that respect volumetric changes, we regularize the cost function using the determinant of the Jacobian of the transformation term to encourage the Jacobian to be close to 1 and prevent the deformation from contracting or expanding unrealistically. The Jacobian can be represented as:

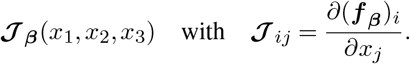

Using the Jacobian, the regularizer is:

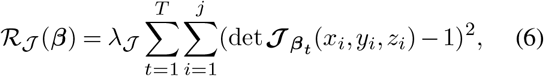

where the Jacobian is evaluated on a grid where we want to ensure its proximity to one.

### 2.5 Optimization

All the variations of the dNMF cost function are optimized in the following way. To update ***β*** and ***A*** we use the `autograd` tool and `PyTorch` library to automatically compute gradients of the cost function and `Adam` optimizer to back-propagate the gradients. A forward pass of computation is evaluating the cost function with ***β***_1:*T*_ and ***A*** (in the fully parametric case, or ***β***_1:*T*_ (in the Gaussian case) as parameters. Note that for a fixed ***C***, all compartments of the cost function are differentiable with respect to the parameters.

To update ***C*** we use multiplicative updates as described in [Taslaman and Nilsson, 2012]:

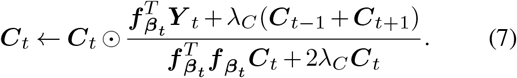

The key difference between these multiplicative updates from those found in [Lee and Seung, 2001] is that the parts of the derivatives of the temporal smoothness regularization terms 2*λ*_*C*_***C***_*t*_ and *λ_C_*(***C***_*t* − 1_ + ***C***_*t*+1_) appear in the denominator and numerator to promote smoothly varying signal.

### 2.6 Initialization

One key advantage of the *C. elegans* datasets considered here is that we can reliably identify the locations of all cells in the field of view, using methods developed in [Yemini et al., 2019]. Using the location of cells in the initial frame (for example) can tremendously aid the optimization of the objective 1 for two main reasons. First, it serves as a very good initializer for the *μ_k_* parameters for cell spatial footprints mentioned in section 2. Second, we know a priori the correct number of cells to be demixed in the FOV. These two factors enable our framework to operate in a **semi-blind** manner towards the deconvolution of neural signals of *C. elegans*, unlike fully blind deconvolution techniques such as e.g. PCA-ICA [Mukamel et al., 2009] or CNMF [Pnevmatikakis et al., 2016].

### 2.7 Using dNMF for image registration

The transformation terms 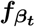, learned using dNMF, can be used to obtain a pixel-level transformation of the video frames to a reference frame in order to yield a registered video; in the ideal case this registered video would remove all the motion from the video, leaving each neuron to flicker in place as its internal activity modulates its fluorescence level. In the current formulation, 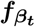 represent push-forward mappings of a reference frame to all the frames in the video. However, to obtain a registration, we need to recover the inverse mappings from all of the video frames to the reference frame. We solve this inverse transform, 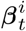, by optimizing the following objective:

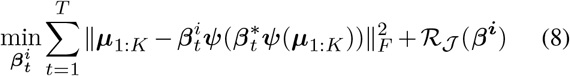

where 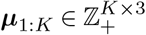 indicates the set of neuron coordinates at the reference frame and 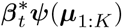 indicates the forward polynomial mapping of these neurons in the *t*-th video frame after optimization, with 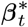 indicating the transformation optimized through Eq. (1). Lastly, 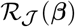 indicates the same Jacobian regularizer as in Eq. (6). In the simplest case that 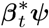 is restricted to be affine, and the regularizer weight *λ_J_* in 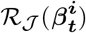 is negligible, then 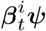 simply implements the shift and matrix inversion of 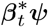. More generally, the exact inverse mapping may not exist or may be unstable; in this more general setting Eq. (8) will output a smooth approximation to the inverse mapping.

Note that Eq. (8) solves a labeled point-set registration problem (since it operates on the neuron centers ***μ***_1:*K*_), not an image registration problem per se. Next we use the recovered inverse mapping 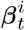 to perform image registration, using pixel-wise interpolation:

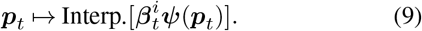

Here, 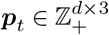 denotes the mesh of pixel coordinates that span the entire volume of the image and Interp. refers to an interpolation function such as linear, nearest neighbor, or bicubic, that can be used to convert non-integer values of pixel co-ordinates to map to discrete pixels. In practice, we set the reference frame to be the first frame in the video series and use linear interpolation. Note that this way of performing registration differs from traditional registration techniques such as Normcorre [Pnevmatikakis and Giovannucci, 2017] in a critical way: the deformation terms that are used to drive the registration are informed by the neural activity and are decoupled from the inferred activity in the joint objective function Eq. (1). Thus, in theory, large fluctuations in neural activity from frame to frame should not affect the deformation terms. In contrast, pure registration techniques on functional neural data may be driven to poor local optima if the neural activity in a particular frame differs strongly from the reference frame.

### 2.8 Population neural analysis

After we have extracted activity traces from each neuron in a single field of view, a typical next step is to compile and analyze a collection of extracted traces across multiple imaged animals. The traces exhibit variability due to both methodological variability (e.g., variability inherent in imaging equipment) and biological variability (e.g., variability inherent in the levels of fluorescent-protein expression across neurons of the same type). These “extra" sources of variability can obscure the changes in neural activity that we wish to extract and analyze here. Consequently, a neuron’s calcium trace, measured across multiple animals, can exhibit differences in overall intensity that require correction to obtain valid comparisons across animals. As a simple example, many neuron classes are composed of a symmetric left and right pair that often show identical calcium activity. With most imaging equipment, when the left neuron is near the lens, the corresponding right neuron is far away, leading to a false differential reading of brightness. Thus, even within a single animal, symmetric neurons can require corrections to be comparable.

The commonly used technique of converting neural traces to Δ*F/F*_0_ aims to correct these issues in mismatched fluorescence intensity profiles but is often insufficient (see Results section below and Figure 2). One way to further normalize time-series data is through z-scoring the signal such that the mean and variance across time is zero and one, respectively. However, in practice, simply mean-shifting to zero often misrepresents the neuron’s baseline signal. Similarly, scaling to unit variance will scale unresponsive and responsive neurons to the same magnitude, thus inflating instead of suppressing measurement noise in unresponsive cells.

**Fig. 2.**
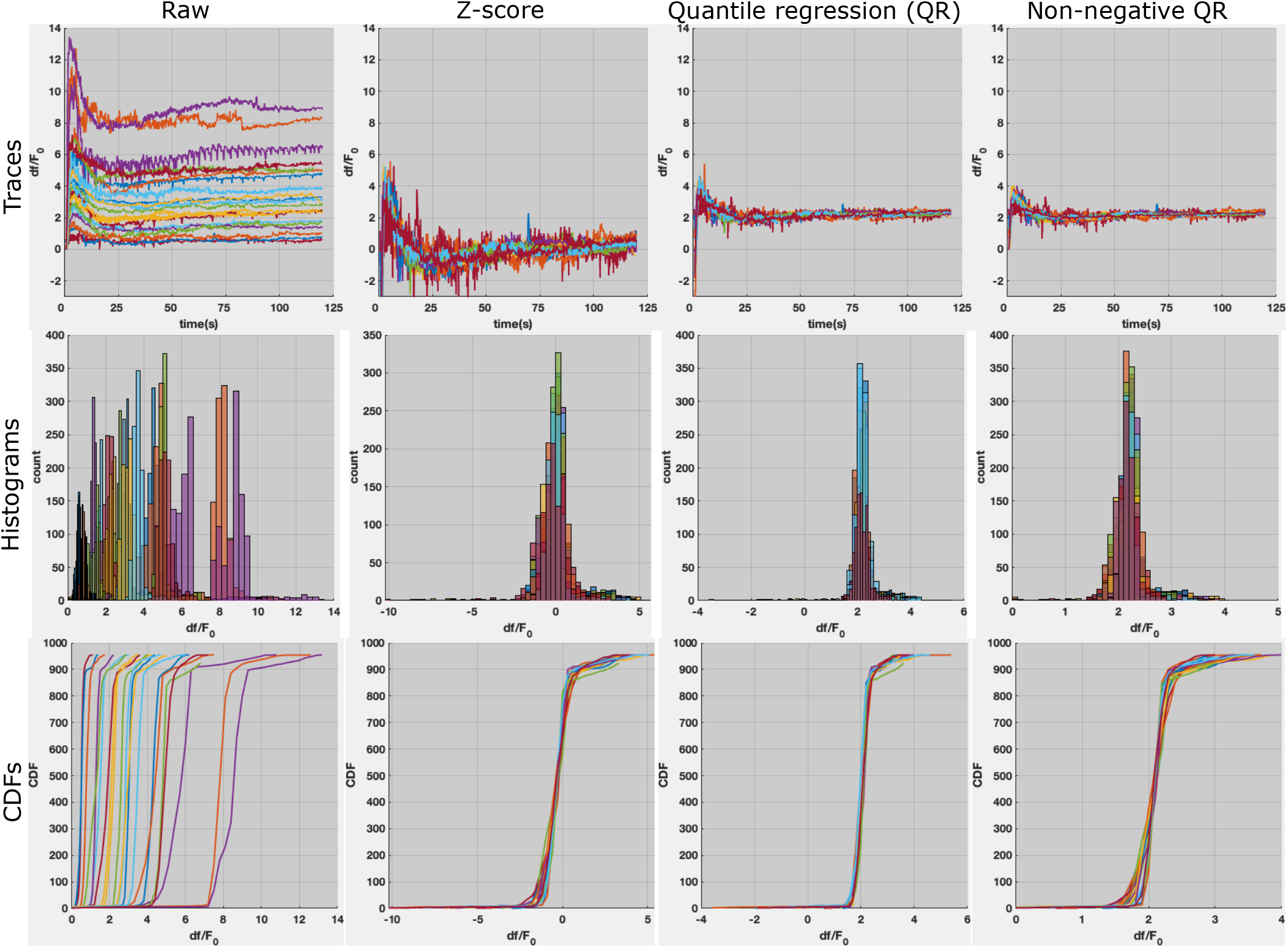
A demonstration of quantile regression for the tail neuron LUAL in *C. elegans*, across 21 animals. Different colors indicate different animals. First column: The raw traces superimposed exhibit variability in intensity profiles due to imaging and biological differences (top). The histograms and cumulative distribution functions (CDFs) of the time-series signals display the differing distributions representing these traces (middle and bottom). **Second column:** Z-scored traces exhibit tighter grouping than raw traces (top) further shown in their CDFs (bottom). However, these z-scored traces are shifted towards zero mean (top) which is misrepresentative of the signal magnitude and also exhibit significant remaining variability. **Third and fourth columns:** After quantile regression (QR) and non-negative quantile regression (NQR) to the medoid of the traces, we see that the normalized traces retain their shape (top), exhibiting even tighter grouping than after z-scoring. In comparison with the z-scored traces, both QR and NQR preserve the median signal magnitude, Δ*F/F*_0_=~2 (top and middle) with smaller tails in their histogram (middle), implying a better fit across the population of animals.

A method that employs a more robust view of the distribution of neural signal would provide a more accurate normalization. Here we generalize the concept of z-scoring time-series by first observing that z-scoring is a linear transform that matches the histogram of the time series to a standard Gaussian distribution with zero mean and unit variance. We then cast histogram normalization in a way such that the transformation is constrained to be a linear transform that minimizes distance to the distribution as a whole, leading to more robust results compared to z-scoring, which restricts attention to two non-robust summary statistics of the histogram (the mean and variance). Lastly, we provide a strategy for normalizing a population of time series data by transforming to the *medoid* of these time series (i.e., the time series which is on average closest to all the others in the population). Empirically, the resulting approach preserves signal while reducing variability across the population.

#### 2.8.1 Quantile regression

Let ***C**^i^* ∈ ℝ^*T*^ denote the time series of a neuron in the *i*th animal over *T* time steps. Suppose we want to match the neural time-series of the *i*th animal to the *j*th animal using a linear transform. One possible strategy to match two time-series signals to one another is to match their baselines and match their peaks. This corresponds to transforming the minimum and the maximum of one time series such that they match the minimum and maximum of the other time series. This is equivalent to matching their minimal and maximal quantiles through a transformation term involving scaling and shifting.

We can generalize this procedure with more quantiles to yield a transformation estimate that is more robust to noise. Matching multiple quantiles using a linear transformation term can be represented by the following linear model:

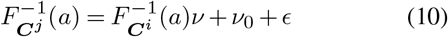

where *F*^−1^ denotes the inverse cumulative distribution function and *ν, ν*_0_ denote the scaling and magnitude shift of the time-series, respectively, and *ϵ* represents an error term. This model posits that each time series signal consists of a baseline and several peaks which can be represented as quantiles of histograms that require matching; baselines and peaks of the same neuron, across different animals, should roughly have similar values.

We can then estimate *ν* and *ν*_0_ by solving the following least squares problem:

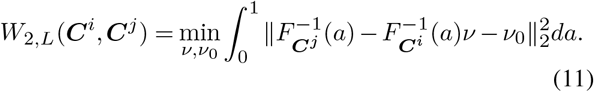

The arg-min of Eq. (11) yields the linear estimates *ν, ν*_0_ that can be used to transform the time series ***C**^i^* to match the time series ***C**^j^*, where 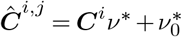. We term this regression model, quantile regression (QR), since the predictors and responses are quantiles of time-series data. If only two quantiles are used i.e. the bottom and topmost quantiles, this procedure is equivalent to matching the minimum/maximum of the two time-series.

Optimizing Eq. (11) yields the transformation that best matches the histogram of the ith time series ***C**^i^* with that of the jth time series ***C**^j^*. The residual discrepancy between the transformed ith time series 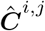 and the *j*th time series ***C**^j^* can be thought of a distance between these time series. In fact, the minimum of Eq. (11) is a linear approximation of a *bona fide* distance metric, termed the Wasserstein metric [Peyré et al., 2019], that is a distance between probability distributions.

Using this notion of proximity between time series, if we have a population of *N* samples ***C***^1^,…, ***C***^*N*^, the strategy for normalizing the time series we advocate here is to compute pairwise Wasserstein distance approximations between all time series and choose the medoid time series to normalize to:

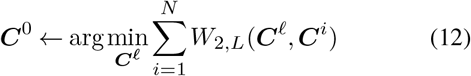

In other words, we can find the best fit of each time series through quantile regression to all other time series, and set as a reference the time series that has the minimal average distance to all the other time series. Once the reference is set, all the samples are transformed to match the reference quantiles using Eq. (11). See Figure 2 for an illustration.

Lastly, if the time series all capture non-negative signal (as is often encountered in calcium imaging) the regression in Eq. (11) can be constrained to be non-negative to ensure the transformed time series maintains its positivity. This yields the non-negative linear estimate of the Wasserstein metric. We term this variant of the quantile regression model as non-negative quantile regression (NQR):

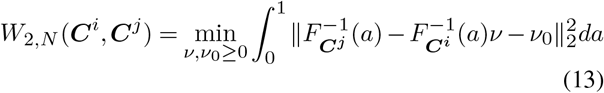

### 2.9 Evaluation metrics

To evaluate the performance of the proposed method as well as the compared methods, we focus on several metrics that shed light both on the signal demixing capabilities of the methods as well as their ability to track objects in time. Namely we focus on two major metrics: **trajectory correlation**, which measures the ability of the deformation model to keep track of the observed motion, and **signal correlation**, which measures the demixing performance by comparing the correlation of demixed signal intensities relative to the ground truth. Specifically, these metrics can be expressed as

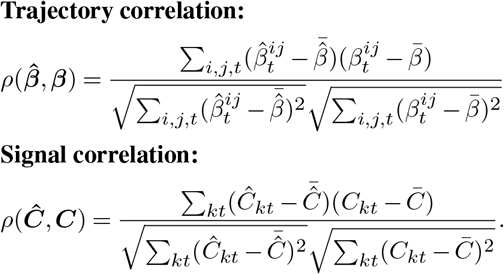

The above metrics are applicable when the ground truth motion trajectories and the signal coefficients are known. To evaluate the performance of the methods using unsupervised registration heuristics, we focus on the **correlation of registered frames** to the average frame (after registration). Heuristically, this measure has been demonstrated to be an effective indicator of successful registration [Pnevmatikakis and Giovannucci, 2017]. Furthermore, in the real data experiments, we also evaluate the average **spread of cell locations** before and after registration. This is computed by taking the average distance of the cells to the average cell location. Similar to the frame correlation measure, this metric allows us to diagnose whether certain cells are registered better than others. While the average correlation of frames is a high-level measure of registration performance, the measure of cellular spread is a localized metric, indicating whether certain regions of the volume are registered better than others. In other words, the former metric measures global sharpness while the latter measures local sharpness.

The added benefit of the latter two evaluation metrics is that since they do not require any ground truth, they can be used for hyperparameter selection; i.e., we can select regularization parameters that yield the sharpest registration results. Furthermore, we can use the sharpness criteria to evaluate the goodness of fit for different deformation models such as quadratic polynomials (as used here), b-splines [Rueckert et al., 1999], or higher order polynomials.

### 2.10 Compared methods

We argue in this paper that jointly optimizing for deformable registration and time-series signal extraction has the potential to improve the quality of both the registration and signal extraction. Therefore, we compare the registration performance of dNMF against the state-of-the-art method for calcium video motion registration, named Normcorre [Pnevmatikakis and Giovannucci, 2017]. Normcorre does not explicitly model the presence of independent signal carrying units in the FOV and instead performs piecewise-rigid transformations on overlapping sub-blocks of the volume using a fast fourier transform based technique [Guizar-Sicairos et al., 2008]. Furthermore, Normcorre uses a normalized cross-correlation registration loss function that is less prone to intensity variations across time-frames.

Next, we also evaluate the signal extraction performance of dNMF against two standard routines in calcium imaging. First, we compare against region of interest (ROI) tracking and pixel averaging within the ROI [Venkatachalam et al., 2016a]. This method tracks the positions of cells across time and extracts signal by taking the average pixel intensity value in a pre-defined radial region around the tracking marker. We also compare against the routine of performing motion correction first and then signal extraction through NMF [Pnevmatikakis et al., 2016]. To replicate this routine in our experiments, we motion correct using Normcorre and then use the Gaussian cell shape parametrization version of NMF that is described in section 2. We use this variant of NMF rather than non-parametric variants such as CNMF [Pnevmatikakis et al., 2016] to bring the comparison against dNMF to an equal footing since dNMF already uses this parametrization that tends to model nuclear shapes well.

### 2.11 Implementation details

All the optimization codes are implemented in `Python 3.7.3` using the autograd tool and the `PyTorch 1.5` package. We used the Adam optimizer with learning rate 0.001 for the simulations and 0.00001 for the worm experiments. Large learning rates lead to jumps in the tracks and lower quality traces, while small learning rates need more iterations to converge. The experiments are run on a Lenovo X1 laptop with Microsoft Windows operating system using 64 GB RAM and Intel(R) Core(TM) i7-8850H CPU @ 2.60GHz, 2592 Mhz, 6 Core(s), 12 Logical Processor(s). We further implemented a sequential optimizer for the demixing of an online stream of videos where each batch of data consisting of a few time frames of the video is processed with parameters initialized using the previous batch. In addition, to improve the memory and time efficiency of our algorithms we also introduced a stochastic variant of dNMF, where in each iteration, to compute the loss and its gradient, we randomly subsample the pixels both in the spatial and temporal domains and update the parameters based on those samples.

### 2.12 *C. elegans* video description

Videos of calcium activity in *C. elegans* were captured via a spinning-disk confocal microscope with resolution (x,y,z)=(0.27,0.27,1.5) microns. Whole-brain calcium activity was measured using the fluorescent sensor GCaMP6s in animals expressing a stereotyped fluorescent color map that permitted class-type identification of every neuron in the worm’s brain (NeuroPAL strain OH16230) [Yemini et al., 2019]. Each video was 4 minutes long and was acquired at approximately 4Hz. Worms were paralytically immobilized (using tertramisole) in a microfluidic chip capable of delivering chemosensory stimuli (salt and two odors) [Chronis et al., 2007, Si et al., 2019]. This setup allows for the controlled delivery of multiple soluble stimuli to the animal with high-temporal precision. See [Yemini et al., 2019] for full experimental details.

Despite paralytic immobilization, we still observed some motion of the worm within the chip, primarily over small distances of several microns and over slow, multi-second time scales. Some of this motion was driven by the animal, while some was the result of the animal drifting passively due to minute pressure differences in the chip. This motion was strongest in the tail, which, due to its taper, was not well secured by the channel walls of the microfluidic chip. Despite the smaller scale of this motion (as compared to freely-moving behavior such as crawling), motion artifacts could strongly confound traces, particularly in the head of the animal where the neurons are very tightly packed. Thus, these motion artifacts required algorithmic correction.

Each dataset from this collection is a video in the form of a 4D tensor *W* × *H* × *D* × *T* (approximately 256 × 128 × 21 × 960) where the value of the tensor at (*x, y, z, t*) corresponds to the activity of a neuron located near the point (*x, y, z*) at time *t*. To extract the neural activity from the videos we first reformat the data into a *d* × *T* matrix where *d* = *WHD* that is called the data matrix ***Y***. We then run the Gaussian dNMF with cell centers initialized using the cell locations in the initial frame, determined using the semi-automated methods described in [Yemini et al., 2019]. Since the cells are approximately spherical in this video we used a fixed spherical covariance matrix for all the cells with squared root diagonal entries equal to 0.57*μ*m (roughly a third of the minimal diameter of adult worm neurons).

### 2.13 Pre-processing steps

Neuron centers were first tracked using a local image registration approach throughout the time series, using the approach in [Venkatachalam et al., 2016a]. After identifying each neuron center in the first frame, every subsequent frame was registered to this first frame. The registration was performed on *x*-, *y*-, and *z*-maximum-intensity projections of a small volume around the neuron center using the imregister function in MATLAB. The volume was chosen to be small enough that nonrigid deformations could generally be neglected, so we used a rigid registration model (translation and rotation only). Because motion is continuous between frames, the initial guess for the transformation was taken to be the calculated transformation from the previous time frame.

We use the initial trajectories of the neurons to initialize our motion parameters ***β**_t_* by solving 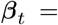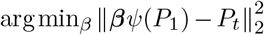 where *P_t_* contains the locations of neurons in time *t* tracked using local image registration techniques. For computational efficiency, we also mask out pixels that are outside of the circles with radii 3*μm* from the location of all neurons in all time points.

## 3 Results

### 3.1 Simulation experiments

To evaluate the effectiveness of our algorithm in capturing motion and demixing time-series traces, we simulated the trajectory of 10 neurons, with a time-specific trace assigned to each (Fig. 3A-B). The signal for each neuron is modeled as a binary vector with length *T* and probability *p* of observing a unit spike, convolved with a decaying exponential kernel. Each trajectory was generated using quadratic transformations of the point cloud in its previous time point, starting from a random initial point cloud. (Note that the composition of many such quadratic mappings is non-quadratic, and therefore the generative model here does not perfectly match the model dNMF uses to fit the data, where a quadratic transformation maps the spatial components *A* to match the observed data at each frame; nonetheless, despite this model mismatch, dNMF achieves accurate results here, as discussed below.) The trajectory of each neuron was then convolved with a fixed 3D Gaussian filter that represents the shape of that neuron and then multiplied with the time course assigned to that neuron. The simulated video is the result of the superposition of these moving Gaussian functions.

**Fig. 3.**
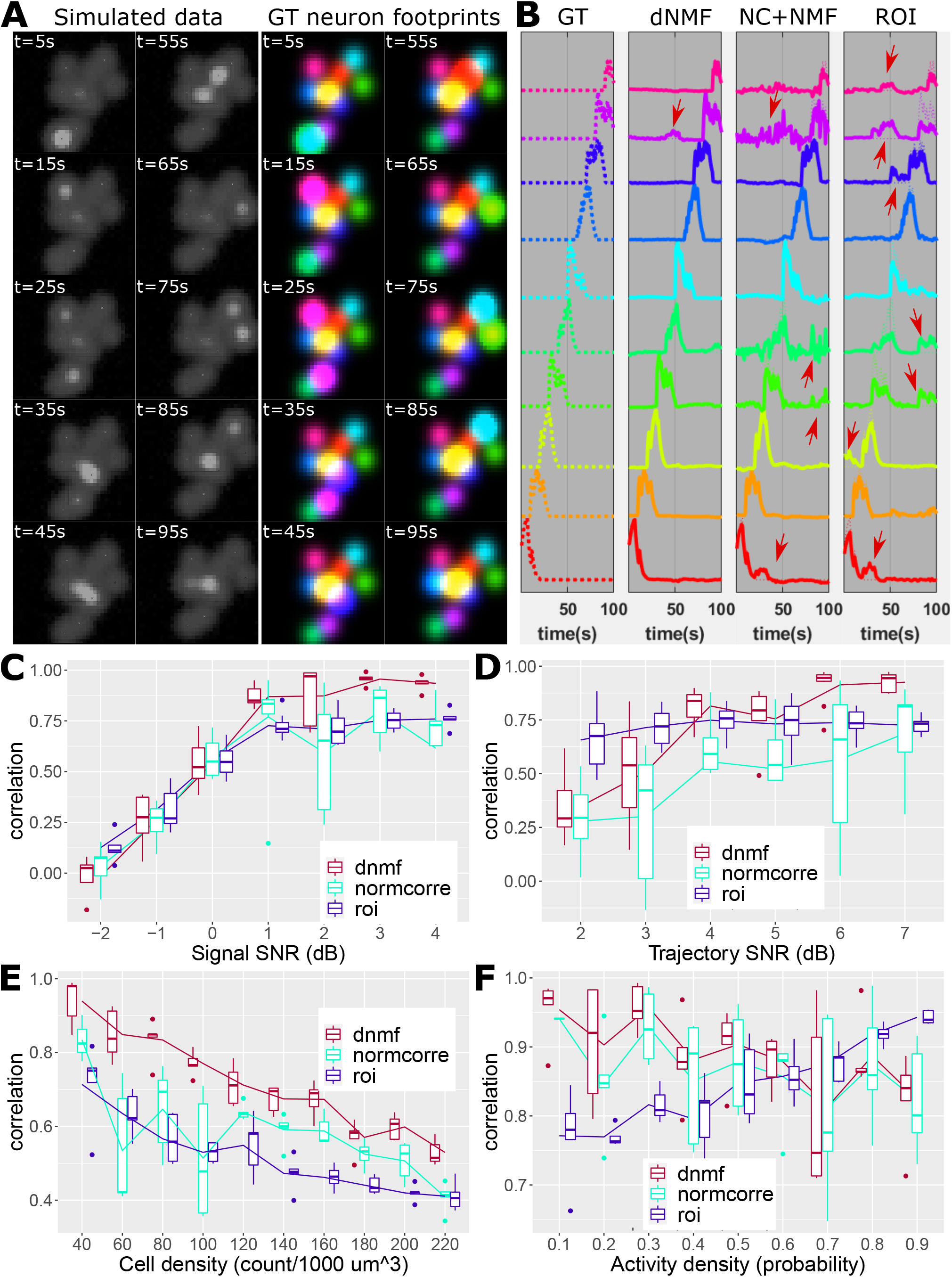
Demixing calcium signals in simulated videos. **A:** Neurons are generated as Gaussian shapes and undergo motion and simulated calcium activity in a 100-second long video. Static snapshots of the video are shown (left) and spatial footprints for each cell are assigned unique colors with intensities proportional to calcium activity (right). Note that the spatial footprints of cells are also in motion, tracking the position of the cells. **B:** The ground truth calcium activity for each cell (left) is compared with the neural activity extracted using dNMF (second column), Normcorre [Pnevmatikakis and Giovannucci, 2017]+NMF (third column) and ROI tracking and pixel averaging (fourth column). dNMF recovers the ground truth signal well whereas Normcorre+NMF and ROI methods yield significantly more mixed signals (indicated by red arrows) due to the proximity of the cells and the tendency of the spatial footprints of mobile cells to overlap. **C:** The correlation of the recovered signal to the ground truth signal as a function of the image signal-to-noise ratio (SNR). **D:** The correlation of the recovered cell movement trajectories to the ground truth trajectories as a function of trajectory SNR. **E:** The correlation of the recovered signals to the ground truth as a function of the density of independent objects in the FOV. **F:** The correlation of the recovered signals to the ground truth as a function of the density of signaling events (simulating neural excitation) exhibited by the cells. Note that we provided ROI tracking here with access to the ground truth cell centers at all times (explaining why ROI averaging correlation values remain high even in the limit of very high activity density); nonetheless, even with artificially perfect tracking accuracy, mixing of nearby signals remains a significant issue. See MOVIE LINK for further details.

We compare the performance of dNMF, Norm-corre+NMF, and ROI pixel averaging in a variety of confounding scenarios using the metrics defined in section 9. In all simulation experiments, the ROI averaging method is provided with the ground truth cell positions — i.e., we examine the accuracy of this method under the (unrealistically optimistic) assumption that neurons are tracked perfectly, to evaluate the demixing performance of ROI signal extraction without the additional confound of tracking performance.

In Fig. 3C-F, we explore the performance limits of dNMF, Normcorre+NMF, and ROI pixel averaging as a function of imaging noise and motion variability. Signal SNR is defined by the peak-to-trough difference between the neural activity signals during times of activity. Trajectory SNR is quantified by how well the cells adhere to the motion of all other cells; high trajectory SNR indicates all cells move in unison, resembling a deformable medium, and low trajectory SNR indicates each cell is moving like independent particles. Mathematically, this is proportional to the log ratio of the variance of the average location of the cells versus the variance of the time differences of these locations. It can be seen in Fig. 3D that dNMF is robust to noise but ultimately may introduce errors to demixing and trajectory tracking if the signal and trajectory SNR (Fig. 3D) are too low. Normcorre+NMF does relatively worse than dNMF as a function of signal SNR and trajectory SNR. ROI pixel averaging has the poorest signal recovery performance of the three compared methods as a function of signal SNR. (Note that ROI pixel averaging enjoys a constant trajectory estimation rate in Fig. 3D, since it has access to ground truth cell locations, as discussed above.)

Next, we evaluated the signal extraction performance as a function of the cell density in the FOV. Increased cell density indicates an increased superpositioning of independent signals and therefore a higher degree of signal mixing. dNMF demixing performance degrades linearly as the density of independent objects within the FOV increases (Fig. 3E) but enjoys higher rates of recovery than both Normcorre+NMF and ROI pixel averaging.

Lastly, we observe that the density of signaling events changes the demixing performance for the three compared methods. In particular, low signal densities (simulating weak excitation) make it harder to track individual cells, which may be dim and therefore hard to detect and track in many frames. dNMF does not suffer in the low signal density regime since it combines information across all visible cells to update the tracking model, and this helps to track dim cells as well.

In Fig. 4, we qualitatively demonstrate the registration performance of dNMF versus Normcorre [Pnevmatikakis and Giovannucci, 2017]. We see that the average frame, after registering with dNMF, is sharper than the non-registered average frame, with better-localized and less-variable cell center locations. In comparison, Normcorre yields a higher spread of cells, even after registration, which may lead to erroneous signal recovery. Both of these global and local sharpness metrics are quantified in Fig. 4C-D.

**Fig. 4.**
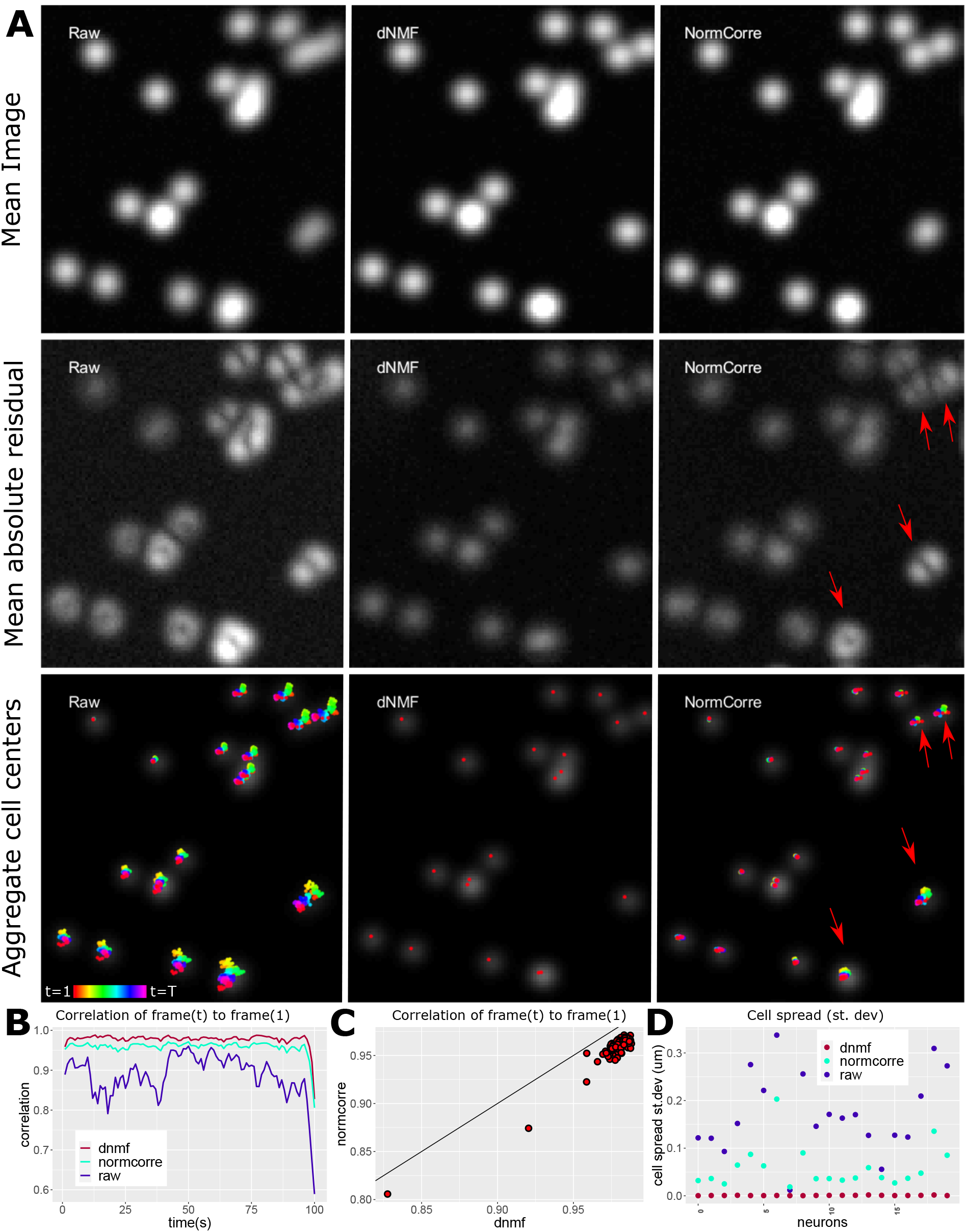
Simulated data registration results. **A: Top row:** The mean video frame prior to registration (left), after dNMF based registration (middle), and after Norm-corre [Pnevmatikakis and Giovannucci, 2017] registration. **Middle Row:** Mean of the absolute value of the video frames subtracted from the first frame prior to registration (left), after dNMF based registration (middle) and after Normcorre (right). If registration is perfect, this image will look like a weighted sum of Gaussian shapes, one for each cell (corresponding to the cell dimming and brightening, but remaining in place); imperfect registrations are indicated by “spreading" or “doubling" of the cell shapes, as indicated by red arrows. **Bottom row:** The locations of the cells across time (colors denote different times) superimposed on the first frame prior to registration (left), after dNMF based registration (middle) and after Normcorre (right). Red arrows indicate cells with imperfect registration, with significant remaining movement of the cells across frames. **B:** The correlation of the video frames to the mean video frame before registration (blue), after dNMF based registration (red), and after Normcorre (cyan); higher values indicate better performance here. **C:** The correlation of individual registered frames to the mean video frame after dNMF registration (x-values) and Normcorre (y-values). The straight line indicates *x* = *y*; points below this line indicate the higher correlation of dNMF registered frames to the mean frame. **D:** The spread of the cell position centers, relative to their average. in the unregistered video (blue), after dNMF-based registration (red), and after Normcorre (cyan). A lower standard deviation for cell spread indicates better performance for local registration of cell shapes. See MOVIE LINK for further details.

### 3.2 Demonstration of demixing in real *C. elegans* data

In the simulated data analyzed above, dNMF exhibits superior registration performance due to its ability to decouple the intensity signal from the motion of objects. Conversely, coupling registration with signal extraction enables dNMF to capture the neural signal and demix it from nearby cells more accurately.

We extend this demonstration further with a real data example. The worm’s tail contains several ganglia, with densely-packed neurons, whose spatial footprints often over-lap due to insufficient spatial resolution. Additionally, even neurons in separate ganglia can end up in sufficient proximity, due to microfluidic confinement or other imaging-setup induced deformations, such that their spatial footprints over-lap. The spatial overlap represents a significant challenge, both for tracking individual neurons and demixing their signal. Figure 5 shows an example of the difficulty present when tracking and demixing neural activity signals from animals with spatially overlapping neural footprints in their recorded images. In this example, ROI tracking loses most of the signal from the LUAR and PLNR neurons and further mixes signals between the DA8/VD13, DVA/DVB, PHCR/PVWL, and PVNL/PVNR neurons. Normcorre+NMF performs better but loses nearly all signals from PLNR while also still mixing signals between the DVA/DVB, PHCR/PVWL, and PVNL/PVNR neurons. In comparison, dNMF recovers strong, independent signals from all ten neurons. Thus dNMF can track and differentiate signals from neurons, even within areas containing multiple spatially-overlapping neural foot-prints where other comparable algorithms fail.

**Fig. 5.**
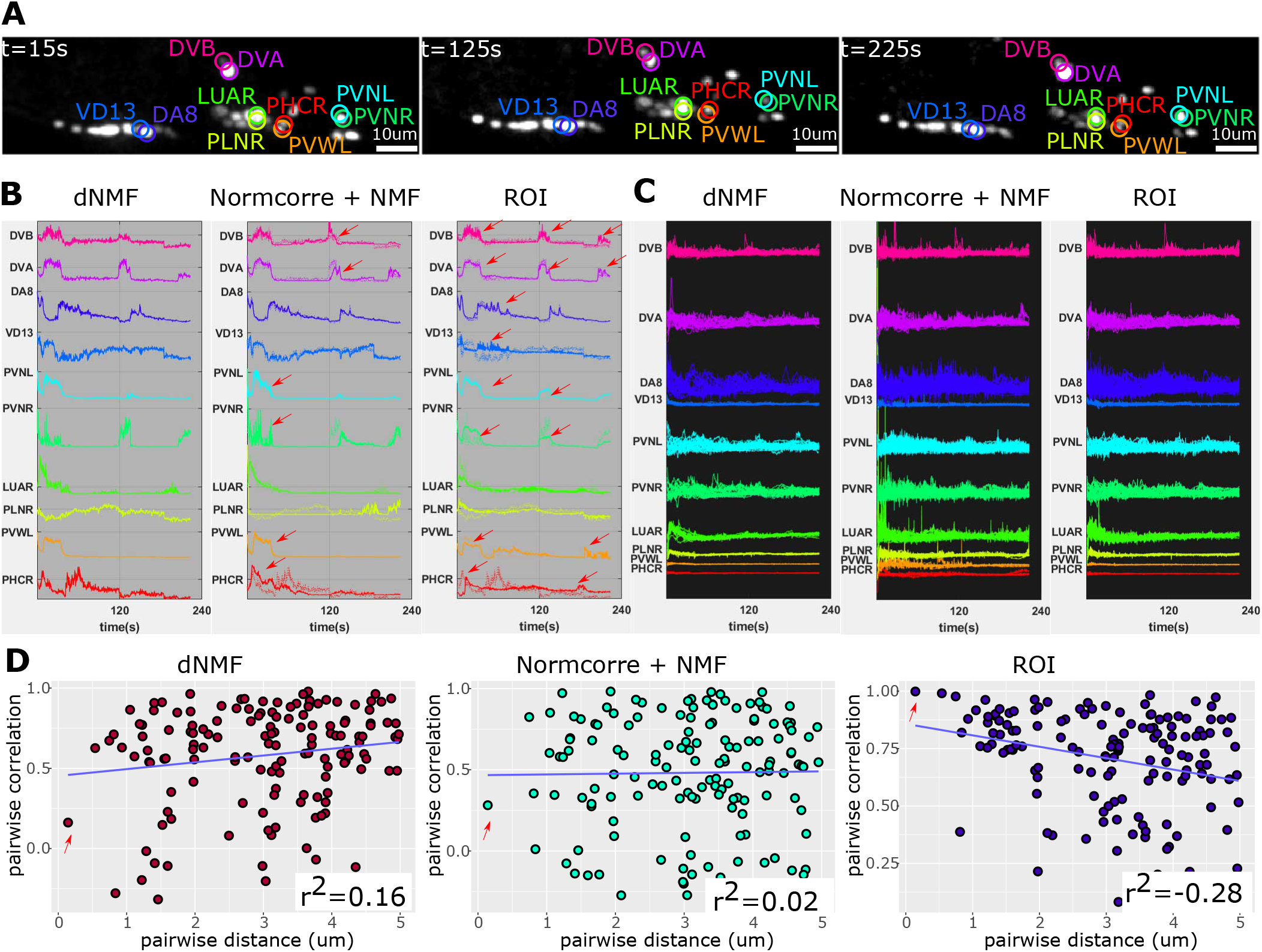
Demixing neural calcium signal in semi-immobilized *C. elegans* videos. **A:** Three static, z-axis maximum projected frames from a representative 4-minute long video of GCaMP6s neural activity. We focus on the signal from five pairs of spatially-neighboring neurons in the tail: DVA/DVB, PVNR/PVNL, PVWL/PHCR, PLNR/LUAR, and VD13/DA8. **B:** Calcium signals extracted by dNMF (left), Normcorre [Pnevmatikakis and Giovannucci, 2017] + NMF (middle), and ROI tracking and averaging (right). dNMF extracts uncoupled signals that demonstrate independent neural activity. The selected cells were chosen such that the signal recovered by ROI averaging is inconsistent with dNMF (quantified by having correlation smaller than 0.4). Normcorre + NMF partially mixes signals between both PHCR/PVWL and PVNL/PVNR around the 30-second mark and DVB/DVA around the 120-second mark (red arrows), and loses nearly all signal from PLNR, due to motion exhibited by the semi-immobilized animal. ROI averaging produces completely correlated signal (red arrows) between all of the labeled neurons, and loses most of the signal from LUAR and PLNR, due to overlap in their spatial footprints. **C:** Calcium activity traces, of the labeled tail neurons, extracted from a population of 21 worms. The unique colors label traces from the same neurons, across different animals. Here, the dNMF traces are tightly grouped, exhibiting minimal variability between animals. Normcorre+NMF traces exhibit mixed-signal and mistracked neurons. ROI traces exhibit wider variability than dNMF, due to mixed signals and, potentially, noise common to ROI averaging. **D:** Pairwise neuron distances versus pairwise correlation of neural signals for all three methods. Note that signal mixing tends to occur when the signal sources are close to one another, necessitating techniques such as NMF to disentangle independent signals. For this reason, dNMF is well suited to demix spatially-close neuron pairs. Normcorre+NMF experiences mixing effects due to motion for which it fails to account (seen in the supplementary movie linked below). ROI averaging does mix traces and thus shows increasingly correlated signals between neuron pairs as they get nearer to each other (indicated by the red arrow). See MOVIE LINK for further details.

Additionally, in figure 5, we quantify the demixing performance by computing the pairwise correlation of nearby neurons as a function of the distance between these neurons. Signal mixing is expected to occur when the spatial footprints of nearby neurons, blurred by the point spread function and/or insufficient spatial resolution, overlap with another. Therefore, one heuristic to determine how well demixing was performed is the correlation of pairwise distances of neurons to the pairwise correlations of their activity. Indeed, both matrix factorization methods, dNMF and Normcorre+NMF, yield uncorrelated trends between neuron pair distance and the correlation of their respective trace activity. On the other hand, simple ROI averaging tends to visibly mix signals in closely-neighboring neurons, resulting in unrealistically high correlation values near 1 for the closest neighbors.

### 3.3 Worm registration

After optimizing dNMF, we can obtain registered videos of worms to evaluate performance and compare against Norm-corre (Figure 6A). Similar to simulated data, we can once again observe that the mean video frame after registering with dNMF is sharper when compared to both the raw average frame and this average after applying Normcorre. Furthermore, the mean of the absolute difference between video frames and the first frame shows that the dNMF registration has fewer distorted toroidal shapes than Normcorre, indicating better registration of cell shapes. Lastly, we can also see that after registering with dNMF the cell centers have a tighter grouping than Normcorre; this is another indication of better registration performance.

**Fig. 6.**
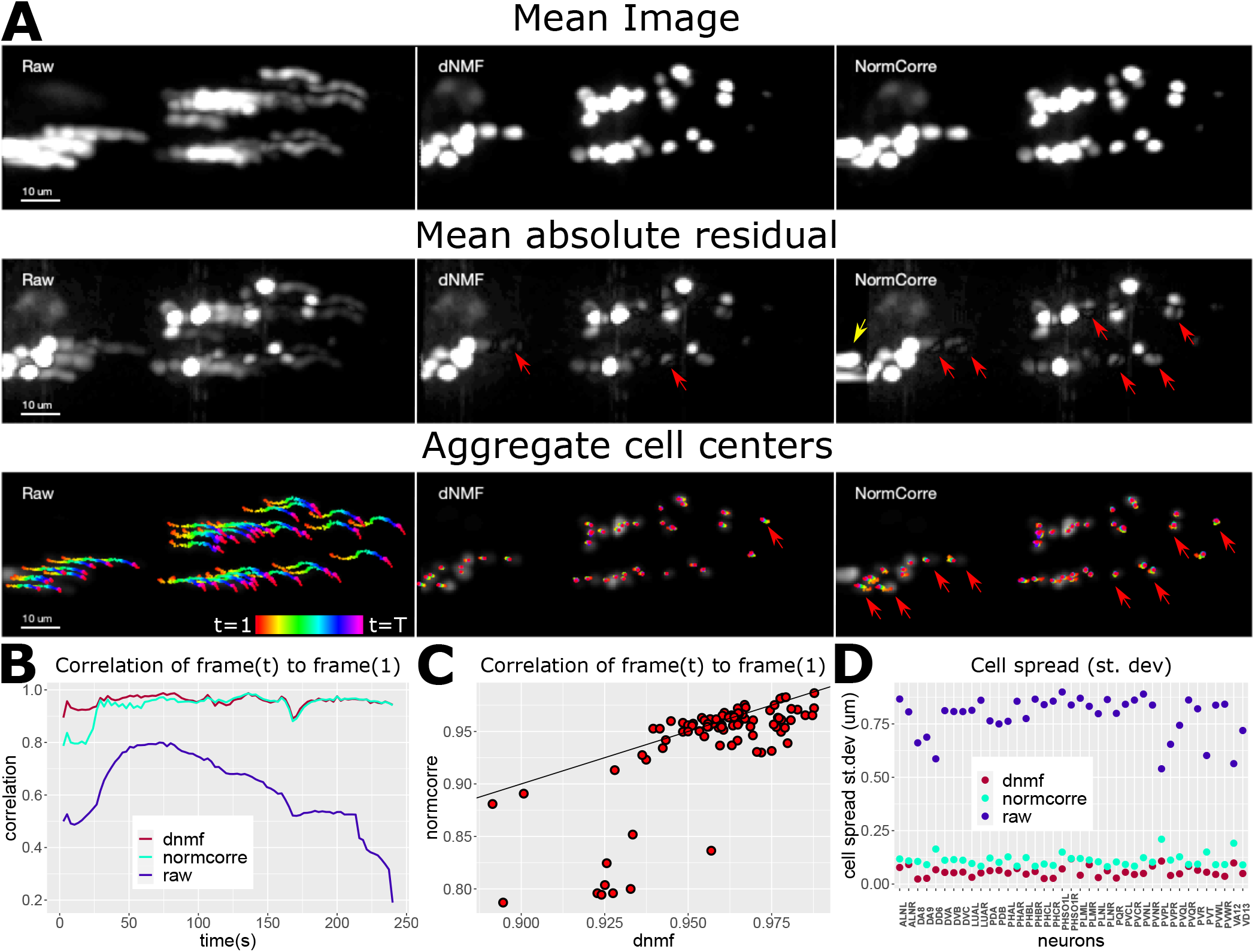
*C. elegans* neural activity video registration results. **A: Top row:** The mean video frame prior to registration (left), after dNMF based registration (middle), and after Normcorre (right). Overall conventions are similar to Fig. 4. **Middle row:** The mean, of the absolute value, of the difference between the first video frame (prior to registration) and subsequent video frames (left). We show these results for dNMF-base registration (middle) and Normcorre (right). Distorted toroidal shapes (indicated by red arrows) denote the superposition of mismatched spatial footprints, indicating a misestimation of deformation. The yellow arrow indicates Normcorre’s boundary pixel extrapolation, which introduces blocky artifacts. **Bottom row:** The positions of cells over time superimposed on the first video frame (left), after dNMF-based registration (middle), and after Normcorre (right). Tighter grouping of cell centers indicates a good correction of motion. Spread groupings of cells indicate poor registration (indicated by red arrows). **B:** Correlation of the video frames to the mean frame, across time, for the unregistered video (blue), after dNMF-based registration (red), and after Normcorre (cyan). dNMF slightly outperforms Normcorre here. **C:** Correlation of the individual video frames, to the mean video frame, after registering with dNMF (x-values) and after Normcorre (y-values). The solid line denotes *x* = *y*. Points below this line (which indicate a higher correlation of registered frames to the average frame) represent better performance for dNMF, and points above this line represent better performance for Normcorre. **D:** The spread of the cell position centers, relative to their average. in the unregistered video (blue), after dNMF-based registration (red), and after Normcorre (cyan). Again, a lower standard deviation for cell spread indicates better performance for local registration of cell shapes. See MOVIE LINK for further details.

Figure 6B-D evaluates these observations quantitatively. The subfigures B and C indicate that the frames registered via dNMF tend to have a higher correlation to the mean registered frame than Normcorre, which indicates the quality of registration. Furthermore, the dNMF and Normcorre registration results diverge most in the initial frames, where the majority of deformable motions are observed. Since Normcorre is a piecewise rigid registration technique, its deformation model may be misspecified to capture such motions, whereas the dNMF motion model is more accurate. Figure 6D demonstrates that the cell grouping after dNMF is indeed tighter than Normcorre.

### 3.4 Population study of *C. elegans*

Using the neural traces extracted with dNMF (converted to Δ*F/F*_0_), we demonstrate the time-series histogram-normalization technique of quantile regression (QR), show its non-negative regression variant (NQR), and compare these with z-score normalization. The time-series data we used is the brain-wide neural GCaMP6s intensity extracted from 21 worm heads (up to 189 neurons in each head) and 21 worm tails (up to 42 neurons in each tail). In these animals, neurons with the same identity often exhibited very different intensity distributions across individual animals. In the course of a time series, neuron intensities change to reflect the underlying activities but, given a sufficiently long recording, after proper alignment, the probability density function (PDF) should be roughly equivalent for neurons of the same class type.

Differences in the intensity PDFs of neurons with identical class types are due to variability in imaging conditions, anisotropy due to random animal orientations, and biological variability in fluorescence expression. To properly compare one animal to another, class-specific neural intensity distributions must be corrected so that they match each other appropriately (e.g., all LUA neurons should exhibit similar PDFs); otherwise, this variability will distort population representations of the signal. In figure 2, we explore these population representations of signal by focusing on a single neuron, LUAL, to compare raw, z-scored, QR, and NQR normalized neural traces. Although the LUAL neurons should preserve similar PDFs, instead they exhibit high variability in both their signal magnitude and baseline activity in their traces, histograms, and cumulative distribution functions (CDFs). Z-scoring partially corrects this variability but retains long tails in the PDFs (histograms), while shifting them to zero mean, which is far less than the median signal observed in the raw traces (a median Δ*F/F*_0_ of approximately 2). In comparison, QR and NQR reduce LUAL neural variability substantially, when compared to z-scoring. Moreover, both QR and NQR preserve the median exhibited by the raw traces and, thereby, retain a better approximation of the neural baseline, whereas z-scoring distorts this baseline.

In figure 7, we extend our demonstration to all head and tail neurons. In this broader representation of neural activity, one can see that the raw traces and even the z-scored traces distort the neural signal, exhibiting a flat appearance with outliers flanking this flattened signal. In contrast, the QR and NQR traces exhibit strong signals without obvious outliers. Thus both QR and NQR can correct variability in neural intensities to help compare signals from neurons with identical types, recorded from a population of animals.

**Fig. 7.**
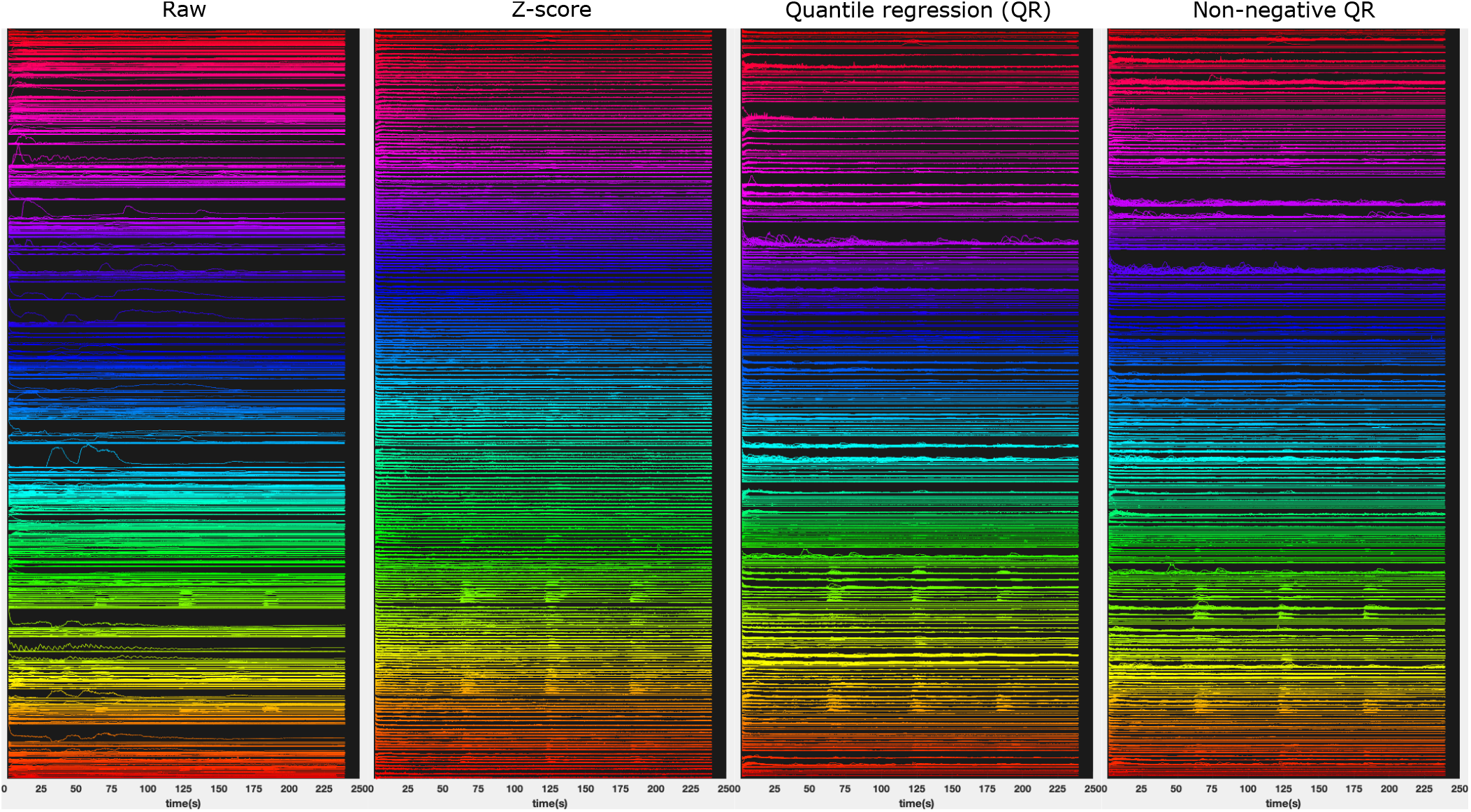
dNMF was used to motion correct, extract, and demix calcium traces of C. elegans neurons from 21 animal heads and 21 animal tails. We demonstrate four strategies for superimposing multi-animal traces for different neuron types. **First column:** Raw traces superimposed, colors indicate different neuron types. Within the same color, different traces indicate different animals. (Y-axis: neuron types), (X-axis) time (s). **Second column:** Z-scored neuron time series. **Third column:** Quantile regression (QR) normalized time series. **Fourth column:** Non-negative quantile regression (NQR). Z-scores use only two summary statistics (mean and variance) for normalization. Z-score scaling to unit variance is strongly influenced by any large-magnitude fluctuations in the signal. Consequently, in a mixture of responsive and unresponsive traces from the same neuron, across multiple animals, z-scored traces with a response will be scaled to match their unresponsive counterparts, thus muting signal in these traces. This is exhibited by the compressed appearance of the z-scored traces in the second column. In contrast, quantile regression uses a more robust and rich set of summary statistics to determine an appropriate scaling. As such responsive and unresponsive neural traces retain appropriate differential scales. This is exhibited by the quantile regression methods shown in third and fourth columns which show better preservation of neuron responses when compared to their z-scored equivalents. Additionally, the z-score translates to zero mean and thus can misrepresent the signal baseline. In contrast, both QR methods preserve the correct signal baseline and, when appropriate, the NQR method can be used to further maintain non-negativity of signals (see Fig. 2).

## 4 Discussion

In this paper, we considered the problem of extracting and demixing calcium signals from microscopy videos of *C. elegans*. We developed an extension of NMF, with a nonlinear motion model applied to the spatial cellular footprints, to deform the static image of these cells, modeling the worm’s posture at each time frame. We provided two different parameterizations for the spatial footprints (Gaussian functions or a non-parametric model) and described regularizations that can help in finding smooth trajectories and signals. We further showed that our method outperforms state-of-the-art models that use a two-step process for motion stabilization/tracking and signal extraction. Finally, we demonstrated the effectiveness of our model by extracting calcium signals from videos of semi-immobilized *C. elegans*.

The dNMF framework we introduce is a generalization of standard non-negative matrix factorization techniques that are commonly used to demix, deconvolve, and denoise neural signals in calcium imaging [Pnevmatikakis et al., 2016]. To tackle the added problem of deforming tissues and mobile animals, we have additionally adopted techniques from the image-registration community to quantify motion in the factorization model. The overall framework is modular and may be further generalized to incorporate different image fidelity loss functions, different regularizers, and alternative deformation parametrizations. Currently, we use a Euclidean loss to assess the goodness-of-fit of the factorization model compared to the observations. Alternative choices for the loss function include normalized local cross correlation [Sotiras et al., 2013]. Similarly, we found it sufficient to use a quadratic polynomial basis for quantifying the motions exhibited by semi-immobilized *C. elegans* worms. However, different animal models or freely-moving worms may necessitate the use of higher-order deformation models such as free-form b-splines [Rueckert et al., 1999].

One common obstacle, after extracting neural traces from a population of animals, is that imaging and biological variability can lead to high variability in the intensity values of traces measured from the same neuron, across multiple animals. To address this, we have provided a time-series normalization technique called quantile regression (QR). This technique bears similarities to z-scoring but it is more robust to outliers in the time series, does not artificially enforce a uniform signal variance over all cells, and yields a more consistent and tighter normalization across a population of neural traces. Moreover, QR can optionally enforce the nonnegativity of the signal that is transformed and normalized (as opposed to z-scoring, which introduces negative values).

Finally, in this work we focused on nuclear-localized calcium imaging in semi-immobilized *C. elegans*. We believe that a similar approach will be useful with other indicators [Chen et al., 2020] and in other preparations, e.g. larval zebrafish [Vanwalleghem et al., 2018], Drosophila [Schaffer et al., 2020], and Hydra [Szymanski and Yuste, 2019]; see, in particular, the recent preprint by [Lagache et al., 2020], who develop improved tracking methods that may nicely complement the dNMF approach. We look forward to exploring these directions further in future work.

## 5 Software

The software is implemented in Python using the Pytorch library and is available at https://github.com/amin-nejat/dNMF.

## Supporting information

Simulation demixing video

Simulation registration video

Worm registration video

Worm demixing video

## Acknowledgements

The authors thank Ruoxi Sun, Conor McGrory, Shreya Saxena, and Scott Linderman for the helpful discussions and suggestions. We also acknowledge the following funding sources. Paninski Lab: NSF NeuroNex Award DBI-1707398, The Gatsby Charitable Foundation, NIBIB R01 EB22913, DMS 1912194, Simons Foundation Collaboration on the Global Brain. Eviatar Yemini: Howard Hughes Medical Institute, NIH (5T32DK7328-37, 5T32DK007328-35, 5T32MH015174-38, and 5T32MH015174-37), Samuel Lab: NIH (1R01NS113119-01) and NSF (IOS-1452593). Albert Lin: NSF Physics of Living Systems Graduate Student Research Network (1806818). Venkatachalam Lab: Burroughs Wellcome Fund Career Award at the Scientific Interface.

## 7 Video captions

### 7.1 Simulation demixing video (VIDEO LINK)

Simulated data demixing results. **Left block:** In the top left we see a max projection of the raw video to be demixed. This video involves 10 neurons in motion with time-varying calcium activity. (For better visibility, note that we use a different simulated dataset here than in Figure 3.) In the middle we see the reconstruction of this video using the moving spatial components extracted with dNMF; each extracted cell is randomly assigned a unique color. On the top right, we see the residual of the raw video minus the reconstruction. The next row shows a selection of cells zoomed and centered on the motion tracking position inferred by dNMF, followed by the corresponding spatial component with intensity proportional to the extracted activity level. Note that the neuron centered panels exhibit little motion, showing that each zoomed cell has been successfully tracked. The following row shows the same cells but with the ground truth signal intensities, and then the localized residual. Finally, the bottom row shows the extracted signals (in the color assigned to each cell) vs. the ground truth (black).

**Right block:** This block has the same conventions as the left block but with Norm-corre [Pnevmatikakis and Giovannucci, 2017] motion corrected video instead of dNMF. The raw data in this panel is motion corrected through Normcorre and thus the reconstruction, residual and the zoom panels are all in the context of the motion corrected video.

As noted above, upon zooming in on the motion corrected patches around each cell, we can see that dNMF provides a good stabilization of the centered cell in each zoomed video, leading to accurately demixed neural signal. In contrast, the Normcorre registered video displays some residual motion which leads to relatively poorer extraction of signal. The high residual signal left by Normcorre+NMF is indicative of poor signal extraction.

### 7.2 Simulation registration video (VIDEO LINK)

Simulated data registration results. **Top row:** The video prior to registration (left), after dNMF based registration (middle), and after Normcorre [Pnevmatikakis and Giovannucci, 2017] registration. **2nd Row:** Absolute value of the video frames subtracted from the first frame prior to registration (left), after dNMF based registration (middle) and after Normcorre (right). If registration is perfect, this frame will look like a weighted sum of Gaussian shapes, one for each cell (corresponding to the cell dimming and brightening, but remaining in place); imperfect registrations are indicated by “spreading" or “doubling" of the cell shapes, as indicated by red arrows. **3rd row:** The estimated deformation field at each time point. **Bottom row:** The locations of the cells across time (colors denote different times) superimposed on the first frame prior to registration (left), after dNMF based registration (middle) and after Normcorre (right). In this video, we see a near perfect stabilization of motion by dNMF, whereas Normcorre displays residual motion as indicated by the absolute residuals (2nd row) as well as the cell spreads in the 4th row.

### 7.3 Worm demixing video (VIDEO LINK)

*C. elegans* video demixing results. **Left block:** In the top left we see the *C. elegans* tail video to be demixed. This video involves about 40 neurons of a semi-immobilized worm tail exhibiting calcium activity. In the right we see the reconstruction of this video using the moving spatial components extracted with dNMF. The top row of the bottom panels show a selection of cells zoomed and centered on the motion tracking position inferred by dNMF and in the bottom row is the corresponding spatial component with intensity proportional to the extracted activity level. Note that the neuron centered panels exhibit little motion, hinting at the successful tracking of objects. **Right block:** This block has the same conventions as the left block but with Normcorre [Pnevmatikakis and Giovannucci, 2017] motion corrected video. **Bottom panel:** This panel shows the raw video with tracking markers superimposed on top of the neuron centers that are used for ROI averaging. In this video, we see that the large motion exhibited by the worm islargely stabilized by dNMF, but less so by Normcorre (as is particularly visible in the zoomed panels).

### 7.4 Worm registration video (VIDEO LINK)

*C. elegans* video registration results. **Top row:** Video prior to registration (left), after dNMF based registration (middle) and after Normcorre [Pnevmatikakis and Giovannucci, 2017] (right) with cell tracking markers superimposed in red. Intensities are oversatured here for better visualization of cell locations. (The flashing blobs in the top left indicate non-neural objects in the FOV such as gut cells.) **Middle row:** The actual intensity profiles of the videos prior to registration (left), after dNMF based registration (middle) and after Normcorre (right). **Bottom row:** The positions of cells over time super-imposed on the first registered frame for unregistered video (left), after dNMF based registration (middle) and Normcorre (right). Tighter cell groups indicate registration that corrects for motion well. Dispersed cell centers indicate residual local motion. In this video, we see that dNMF has slightly better registration performance than Normcorre as exhibited by the smaller cell spread in several neurons, especially the neurons in the pre-anal ganglion (towards the left border of the frame).

## 8 Appendix

## 8.1 Spatial component: non-parametric model (additional details)

Similar to the standard NMF models, we can parameterize ***A*** using an *d* by *k* matrix where *d* is the number of pixels of one time frame of the video and *k* is the number of objects that are present. 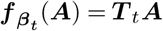 where ***T**_t_*: ℝ^*d*×*d*^ and

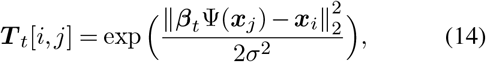

where ***β**_t_* is a 3 by 10 matrix, and Ψ: ℝ^3^ → ℝ^10^ is the polynomial basis function that maps a 3D location into its 10-dimensional quadratic representation in the following way Ψ([*x, y, z*]^*T*^) = [1*, x, y, z, x*^2^, *y*^2^, *z*^2^*, xy, yz, zx*]^*T*^. The choice of *σ* controls the amount of the spread of the mass of a pixel into nearby pixels. If *σ* is large then each pixel will diffuse into many pixels around it which will create artificial overlap between the component footprints. If *σ* is very small then we get vanishing gradients. A careful choice of *σ* is therefore important for the optimization. We usually set the *σ* to 0.1 in our experiments. If we set the ***β***_1:*T*_ and ***T***_1:*T*_ to the identity we will recover the original formulation of NMF [Lee and Seung, 2001]. However, if we optimize over ***β***_1:*T*_ we can capture the nonlinear deformation of the objects and estimate the time varying signals more accurately. Furthermore, using this type of kernel matrix as a surrogate for the quadratic transformation has the advantage that it allows us to compute gradients of the cost function with respect to ***β**_t_*:

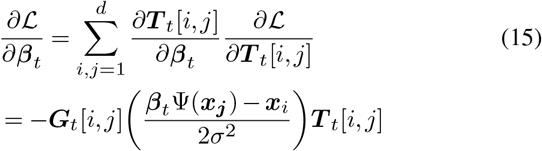

where 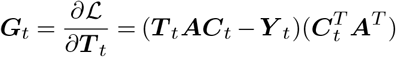

We can also compute the gradient with respect to other parameters of the model similar to the vanilla NMF and derive the optimal step size for faster multiplicative updates:

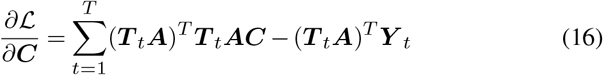

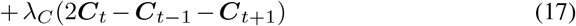

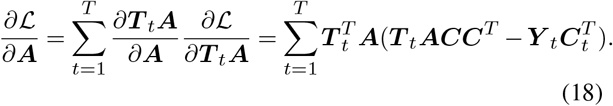

## 8.2 Regularization: spatial location priors

One of the key advantages of our modular framework is that, if available, we can use statistical atlases of neural positions to better condition our non-convex objective function towards favorable local minima. This is especially useful in the non-parametric modeling of spatial components, where we may not have a good model of the spatial occupancy map of cells.

Similar to the approach taken by [Saxena et al., 2019], we can introduce a regularizer to encourage spatial proximity of non-parametric components to a set of predefined centers that is defined by the statistical atlas of neuron positions. Given a (non-negative) constraint matrix ***D*** ∈ ℝ^*d*×*k*^ such that each column encodes the allowable occupancy maps of each of the *k* cells, the regularizer is:

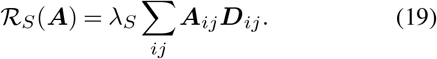

The constraint matrix ***D*** determines the span by which pixels can be part of a spatial footprint for each component. For example, the *k*th column of ***D*** has 1’s in the entries that correspond to plausible pixel locations that the *k*th neuron may occupy. Alternatively, ***D*** can contain weighted values such that pixels with smaller distance values to a component will tend to have a larger weights.

The utilization of this spatial location regularization using statistical atlases (along with the corresponding strong prior information on the number of cells that should be visible in the field of view) is an important advantage of dNMF (and loca-NMF [Saxena et al., 2019]) over blind deconvolution techniques such as CNMF [Pnevmatikakis et al., 2016].

